# Low alcohol preferring mice have reduced task engagement during a waiting task for alcohol, which is enhanced by intermittent alcohol drinking

**DOI:** 10.1101/2022.05.25.493462

**Authors:** Phillip Starski, Danielle Maulucci, Hunter Mead, Frederic Hopf

## Abstract

Alcohol use disorder (AUD) is related to excessive binge alcohol consumption, and there is considerable interest in associated factors that promote intake. AUD has many behavioral facets that enhance inflexibility toward alcohol consumption, including impulsivity, motivation, and attention. Thus, it is important to understand how these factors might promote responding for alcohol and can change after protracted alcohol intake. Previous studies have explored such behavioral factors using responding for sugar in the 5-Choice Serial Reaction Time Task (5-CSRTT), which allows careful separation of impulsivity, attention, and motivation. Importantly, our studies uniquely focus on using alcohol as the reward throughout training and testing sessions, which is critical for beginning to answer central questions relating to behavioral engagement for alcohol. Alcohol preference and consumption in C57BL/6 mice were determined from the first 9 sessions of 2-hour alcohol drinking which were interspersed among 5-CSRTT training. Interestingly, alcohol preference but not consumption level significantly predicted 5-CSRTT responding for alcohol. In contrast, responding for strawberry milk was not related to alcohol preference. Moreover, high-preference (HP) mice made more correct alcohol-directed responses than low-preference (LP) during the first half of each session and had more longer reward latencies in the second half, with no differences when performing for strawberry milk, suggesting that HP motivation for alcohol may reflect “front-loading.” Mice were then exposed to an Intermittent Access to alcohol paradigm and retested in 5-CSRTT. While both HP and LP mice increased 5-CSRTT responding for alcohol, but not strawberry milk, LP performance rose to HP levels, with a greater change in correct and premature responding in LP versus HP. Overall, this study provides three significant findings: 1) alcohol was a suitable reward in the 5-CSRTT, allowing dissection of impulsivity, attention, and motivation in relation to alcohol drinking, 2) alcohol preference was a more sensitive indicator of mouse 5-CSRTT performance than consumption, and 3) chronic alcohol drinking promoted behavioral engagement with alcohol, especially for individuals with less initial engagement.

## 1. INTRODUCTION

Excessive alcohol consumption is a prevalent activity that may progress to Alcohol Use Disorder (AUD), and ∼3/4^th^ the ∼$250 billion/year cost of drinking in the US comes from the ∼1/7^th^ of adults who binge (1). Excessive intake can contribute strongly to the substantial harms of alcohol, including enhanced risk of drinking problems (2-4), while reducing excess intake lowers health risks and relapse (5-7). Higher risk for binge drinking has been linked to high trait impulsivity (8-10), and non-dependent drinkers with higher self-reported impulsive behavior achieve higher blood alcohol levels during free-access self-administration, and experience greater euphoria from alcohol (11). Impulsivity is complex construct (12-14), with variants related to motor (impulsive action) and cognitive (impulsive choice) functions (see Discussion), and is considered an important risk factor for AUD. As this disorder develops, it is accompanied by significant changes in cognitive behavioral control (15, 16). The desire for intoxication and the increased tolerance of adverse consequences are examples of motivational changes in people with AUD (17-19). Further, an “attentional bias” will typically develop that promotes behavior toward alcohol cues over natural rewards (20-22). Together, impulsivity, motivation, and attention are key aspects of behavioral engagement with alcohol that we seek to investigate, especially changes in such responding after chronic alcohol use.

The five-choice serial reaction time task (5-CSRTT) is a multifaceted behavioral paradigm that has been thoroughly characterized in rodents to elucidate impulsive, attentional, motivational, and perseverative behavior in the same session (23-25). Thus, the 5-CSRTT is valuable for assessing a broad range of measures of behavioral performance, when compared to many other tasks. Interestingly, a 5-CSRTT version adapted for humans predicts higher alcohol intake in more impulsive individuals, suggesting high translational value (26). While determining clear correlations between behavioral factors in human studies remains challenging, rodent studies give the ability to dissect important contributors to behavioral engagement for alcohol. However, to date, studies examining the relation of alcohol and impulsivity have primarily determined 5-CSRTT responding for sugar in relation to alcohol exposure (25-31).

Here, we have uniquely adopted the 5-CSRTT to have alcohol as the reward, allowing us greater precision in identifying the nature of behavioral engagement, with the goal of understanding how impulsivity, motivation, and attention for alcohol might relate to preference or consumption. Interestingly, we found that 5-CSRTT performance was significantly related to alcohol preference rather than consumption level. Thus, it is interesting that, in addition to high trait impulsivity, people at risk for binging have higher alcohol preference (8-10), and impulsivity can be linked to preferences in rodents (32-37) (see Discussion). In addition, after chronic alcohol consumption, mice overall increased their performance, but this was especially pronounced in initially low-responding mice, suggesting that protracted drinking may be particularly hazardous for individuals with lower initial drive for alcohol. Finally, we also performed several days of 5-CSRTT responding for strawberry milk, with our previously used methods (27, 28). Sweet milk responding was higher than alcohol and with greater accuracy, did not relate to alcohol preference, and had minimal changes with chronic drinking, suggesting important specificity in the alcohol-engagement relationship. We provide herein a robust and valuable model to help understand inter-relationships between different aspects of engagement for alcohol, and how they could be altered by chronic drinking, which together promote excessive intake.

## 2 MATERIALS AND METHODS

### 2.1 Animals

Forty-eight male C57BL/6J mice from Jackson Laboratories Inc were individually housed, starting at 8 weeks old, in standard Plexiglass cages with *ad libitum* access to food and water until water restriction. Mice were maintained in a 12h:12h reverse light-dark cycle. Animal care and handling procedures were approved by the Indiana University Institutional Animal Care and Use Committee in accordance with NIH guidelines.

### 2.2 5-Choice serial reaction time task (5-CSRTT)

A detailed description of early-stage, late-stage, and strawberry milk (SM) training can be found in supplementary methods. All mice were trained and tested under a 10s stimulus duration (SD) and 5s intertrial interval (ITI): this was done to reduce challenge within the task, since this is, to our knowledge, the first investigation using an intoxicant (10% alcohol) as the reward in 5-CSRTT. Figure 1A shows the overall timeline of studies, and Figure 1B (see figure legend) and Figure S1 give visual representations of several typical session events.

**Figure 1.**
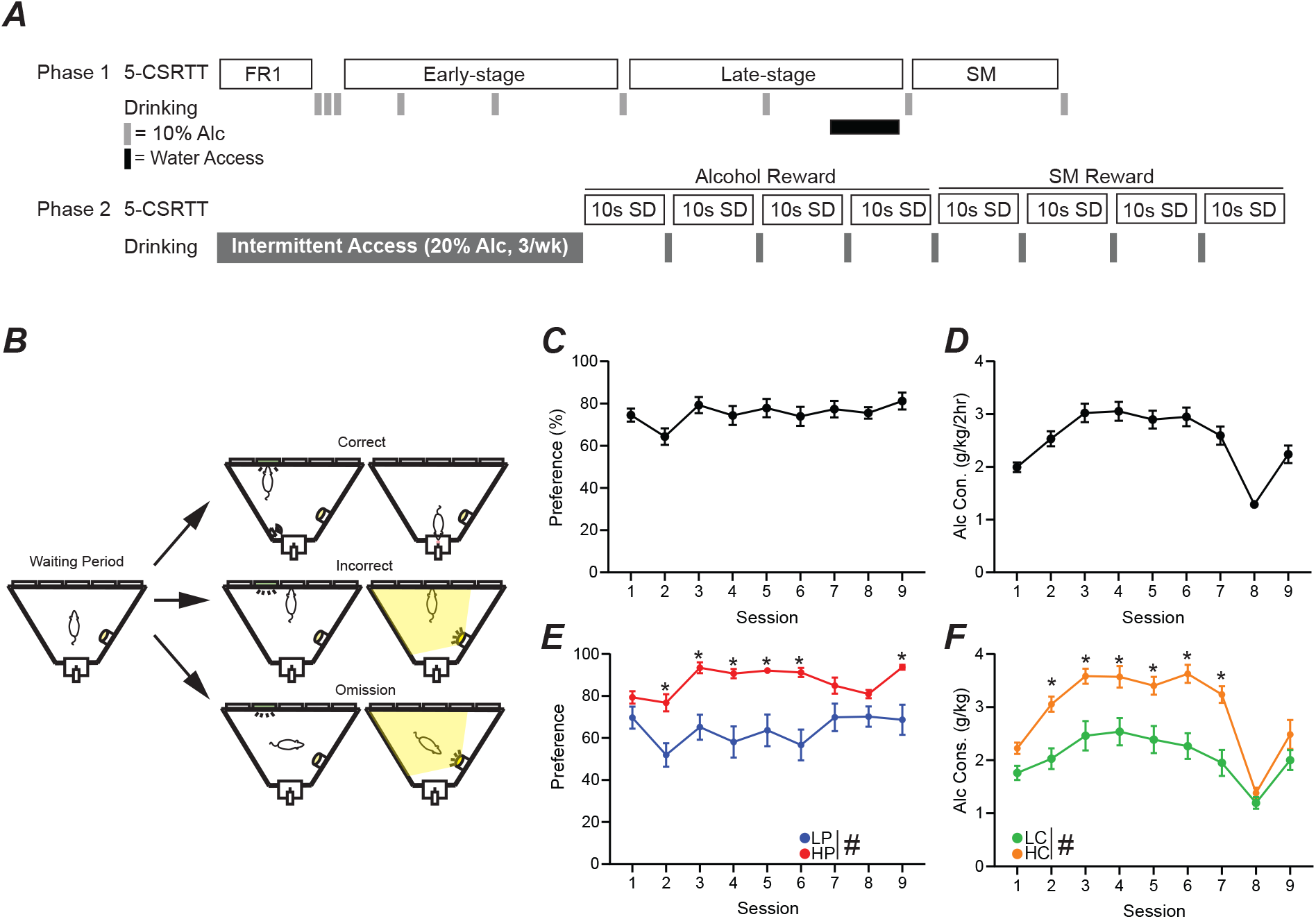
5-Choice Serial Reaction Time Task training schedule and alcohol drinking. (A) Schedule of 5-CSRTT and drinking behavior. (B) Graphic detailing a correct, incorrect, and omission within the 5-CSRTT. Briefly, a head entrance into an illuminated reward tray initiates the waiting period. When the waiting period ends, a light will appear in 1 of 5 ports. A touch in the illuminated port is a correct response and will result in reward delivery. A touch in an unlit port is an incorrect response and will cause a punishing flash of light and no reward. If the lit port extinguishes and the limited hold period elapses, this will result in an omission causing a punishing flash and no reward. Overall (C) alcohol preference and (D) alcohol consumption of all mice, across the 9 days of 2-hour DID two-bottle choice drinking (gray bars). (E) Alcohol preference when mice were separated by median split of preference (HPvLP: F_1,46_=82.06, *p*<0.0001). Post-hoc revealed significance on sessions 2-6 and 9. (F) Alcohol consumption when mice were separated by median split of consumption (F_1,46_=81.31, *p*<0.0001; time: F_6.11,279.4_=18.30, *p*<0.0001; interaction: F_8,366_=2.22, *p*=0.0253). Post-hoc revealed significance on sessions 2-7. Session 8 reflects the time in which the mice were given one week of water access before being water restricted again. #*p*<0.05 for group effect. \**p*<0.05 post-hoc significance. All data are expressed as ±standard error mean.

2-hour drinking-in-the-dark (DID) sessions, with access to a bottle of 10% alcohol versus a bottle of water (based on our previous method in Lei et al. 2019 (38), were interspersed among 5-CSRTT test sessions (as shown in Fig.1) to assess drinking preference and level within each mouse.

### 2.3 Intermittent Access

The intermittent access two-bottle choice (IA2BC) paradigm for 20% alcohol was based on Lei et al. 2019 (38) and are detailed in the supplementary material.

### 2.4 Statistical Analysis

5-CSRTT and alcohol consumption studies were analyzed by two-way repeated measures analysis of variance (ANOVA) followed by Bonferroni’s multiple comparisons test where appropriate. Sphericity was not assumed, and the Geisser-Greenhouse correction was used. For missing data points (spill during drinking, or animal non-responding), a mixed-model analysis of variance was used. All group analyses of pre-IA2BC versus post-IA2BC and performance changes were tested for normal distribution (Shapiro-Wilk test), and then an appropriate test (parametric or non-parametric) was used to measure differences (t-test, Mann-Whitney, Paired t-test, Wilcoxon). Non-parametric data is reported simply in the main text, with specific values in the supplementary material. All statistical analyses were calculated using Prism 9.0 software (Graphpad Software Inc., San Diego, CA), with significance set at *p*<0.05.

## 3 RESULTS

### 3.1. Mice with high alcohol preference or consumption show higher engagement in early-stage training

To better understand the relationship between alcohol drinking behavior and 5-CSRTT performance, drinking preference and consumption was calculated from the first nine 2 hour-DID two-bottle-choice sessions (Figure 1C,D). A median split was then performed to compare response patterns in High alcohol Preference (HP, n=24) versus Low Preference (LP) mice (n=24, Figure 1E), and a separate median split was performed to compare high consumption (HC, n=24) versus low consumption (LC, n=24) mice (Figure 1F). For all subsequent analyses, performance measures were analyzed separately by preference and by consumption.

Each 5-CSRTT session involved unlimited trials in an hour period, where a nosepoke to an illuminated stimulus in one of five ports led to rear reward delivery (detailed further in Figure 1B legend). During the first ten days of training (“early-stage”), HP mice performed significantly more trials than LP mice (Figure 2A_1_, HPvLP: F_1,46_=11.36, *p*=0.0015; time: F_1.93,88.74_=9.59, *p*=0.0002), with higher accuracy (Figure 2A_2_, HPvLP: F_1,46_=4.84, *p*=0.0329; time: F_4.81,221.2_=6.57, *p*<0.0001; interaction: F_9,414_=1.96, *p*=0.0421) and significantly more correct responses compared to LP mice (Figure 2A_3_, HPvLP: F_1,46_=11.16, *p*=0.0017, time: F_1.76,80.8_=8.5, *p*=0.0008; interaction: F_9,414_=4.57, *p*<0.0001). In addition, the percent of trials that were omissions in early-stage was lower in HP mice (Figure 2A_4_, HPvLP: F_1,46_=11.4, *p*=0.0015; interaction F_9,414_ = 2.48, *p*=0.0091). However, raw premature responses were significantly higher in HP mice (Figure 2A_5_, HPvLP: F_1,46_=6.96, *p*=0.0113; time: F_1.39,64.07_=4.13, *p*=0.0334; interaction: F_9,414_=2.48, *p*=0.0091), as were the percentage of premature responses (Figure S3G, HPvLP: F_1,46_=7.04, *p*=0.0109; time: F_2.71,124.6_=2.93, *p*=0.0411; interaction: F_9,414_ = 3.27, *p*=0.0007). Together, these suggest that HP have greater engagement than LP mice early in training, with greater number of trials, correct, and accuracy, and also fewer omissions and greater premature responding (which could reflect greater impulsivity, or simply greater engagement with the task).

**Figure 2.**
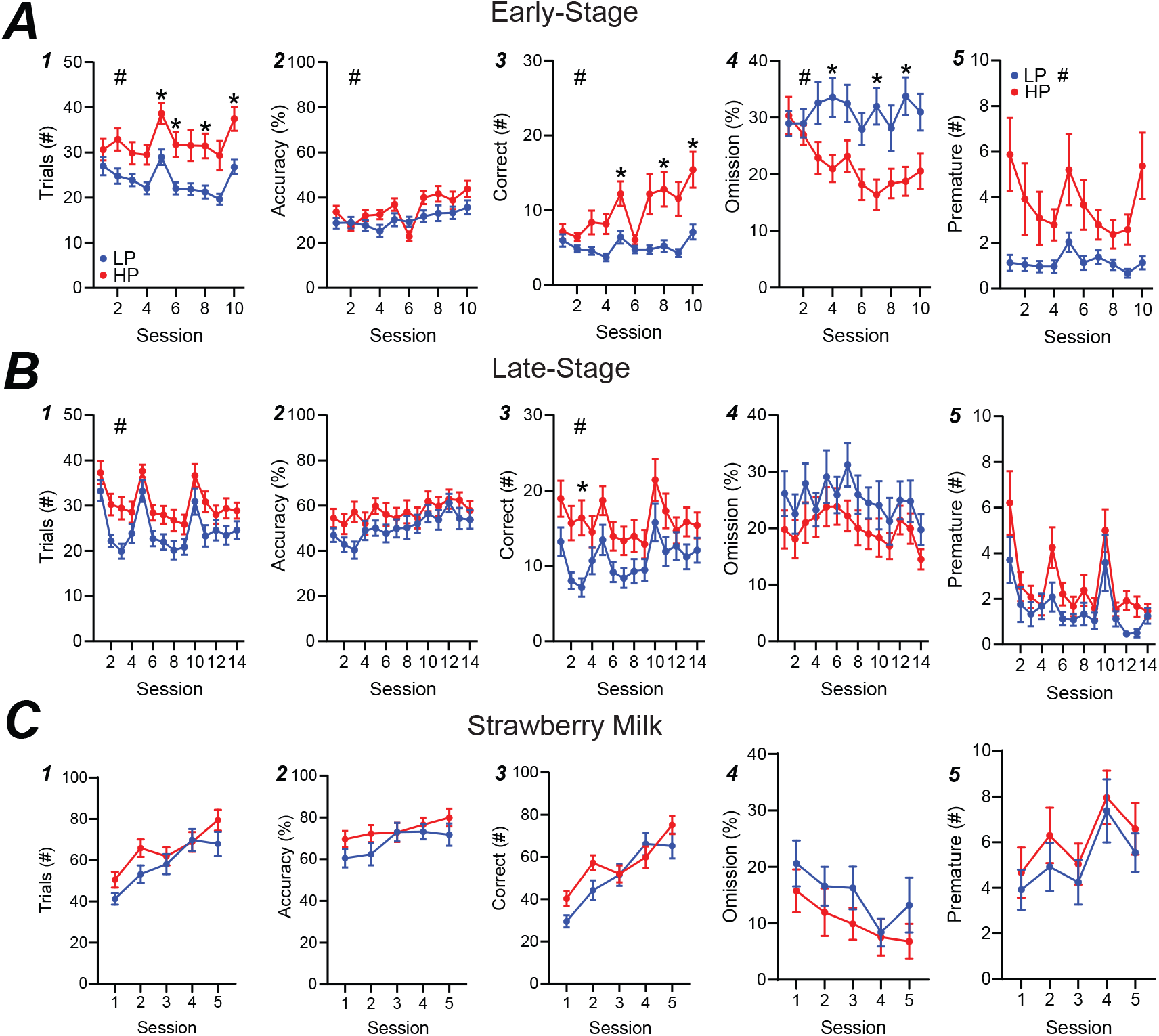
HP versus LP early-stage and late-stage alcohol, and SM training in the 5-CSRTT. (A) Early-stage performance showed differences in trials, accuracy, correct (post-hoc significance on session 3), %omission, and premature between HP and LP mice. (B) Late-stage performance displayed HP-LP differences in trials and correct, but not accuracy, %omission or premature. (C) SM performance showed no differences in any metric. n= 24/group for preference. #*p*<0.05 for group effect. \**p*<0.05 post-hoc significance. All data are expressed as ±standard error mean.

Similar to our HP versus LP comparison, we found that mice with HC overall performed more in early training trials than LC mice (Figure S2A_1_, HCvLC: F_1,46_=4.12, *p*=0.0482; time: F_1.81,83.1_=9.56, *p*=0.0003) and had more correct responses than LC mice (Figure S2A_3_, HCvLC: F_1,46_=4.57, *p*=0.0380; time: F_1.67,76.91_=8.11, *p*=0.0013; interaction: F_9,414_=2.25, *p*=0.0185), although with no post-hoc differences in any session. In addition, and unlike HP versus LP, HC and LC mice were not different in accuracy (Figure S2A_2_, HCvLC: F_1,46_ = 1.34, *p*=0.2528; time: F_4.55,209.3_=6.47, *p*<0.0001) or percentage of omissions (Figure S2A_4_, HCvLC: F_1,46_=3.44, *p*=0.0699). Together, these suggest that differences in performance early in training were more related to alcohol preference differences rather than consumption.

### 3.2 Preference predicts performance, while alcohol consumption does not, during late-stage training

In late-stage training, we continued with a 10s SD and 5s ITI, since we wanted to explore potential behavioral differences under simpler task requirements in this first-time assessment of 5-CSRTT with alcohol as the reward. HP and LP mice continued to show significant performance differences in late-stage training, while HC and LC mice did not. HP mice performed more trials (Figure 2B_1_, HPvLP: F_1,46_=5.93, *p*=0.0188; time: F_7.74,355.9_=18.91, *p<*0.0001) and correct responses (Figure 2B_3_, HPvLP: F_1,46_=4.84, *p*=0.0329; time: F_7.2,331.4_=9.11, *p*<0.0001) than LP mice. However, by late-stage training, there were no differences between preference groups for accuracy (Figure 2B_2_, HPvLP: F_1,46_=1.92, *p*=0.1730; time: F_7.52,345.8_=5.37, *p*<0.0001), omissions (Figure 2B_4_, HPvLP: F_1,46_=1.88, *p=*0.1772; time: F_6.81,313.1_=3.01, *p*=0.0049), raw premature responses (Figure 2B_5_, HPvLP: F_1,46_=3.801, *p*=0.0573; time: F_4.90,225.4_=10.48, *p*<0.0001), or percentage of premature responses (Figure S3H, HPvLP: F_1,46_=3.96, *p*=0.0525; time: F_8.71,400.5_=5.43, *p*<0.0001). In contrast, HC and LC mice did not show differences in trials performed (Figure S2B_1_, HCvLC: F_1,46_=1.71, *p*=0.1979; time: F_7.8,359_=19.10, *p*<0.0001), accuracy (Figure S2B_2_, HCvLC: F_1,46_=0.119, *p*=0.7319; time: F_7.56,347.8_=5.34, *p*<0.0001), number of correct responses (Figure S2B_3_, HCvLC: F_1,46_=0.717, *p*=0.4016; time: F_7.45,342.9_=9.2, *p*<0.0001; interaction: F_13,598_=1.89, *p*=0.0286), or omissions (Figure S2B_4_, HCvLC: F_1,46_=0.139, *p*=0.7115; time: F_6.74,309.9_=3.03, *p*=0.0048). Together, our data suggest, perhaps surprisingly, that preference is a better indicator of established (later-stage) performance under a more basic version of 5-CSRTT than consumption, since HP had more trials and correct responses than LP, while HC and LC were not different.

### 3.3 Unlike alcohol, performance for strawberry milk reward is not related to alcohol preference

After late-stage alcohol testing, mice were switched for 5 sessions to a strawberry milk (SM) reward in the 5-CSRTT to determine whether preference-related performance for alcohol (Figure 2A,B) might be related to more basic differences in motivation for reward learning. Overall, mice had more than twice the number of responses for SM relative to alcohol (Figure S4M, t[94]=10.38, *p*<0.0001). However, there were no differences in any response measure between HP and LP mice, including in number of trials (Figure 2C_1_, HPvLP: F_1,46_=1.571, *p*=0.2163; time; F_2.76,127.0_=42.46, *p*<0.0001; interaction: F_4,184_=3.017, *p*=0.0193), accuracy (Figure 2C_2_, HPvLP: F_1,46_=1.278, *p*=0.2639; time: F_2.81,129.3_=7.62, *p*=0.0001), correct responses (Figure 2C_3_, HPvLP: F_1,46_=1.242, *p*=0.2709; time F_2.73,125.5_=43.63, *p*<0.0001), omissions (Figure 2C_4_, HPvLP: F_1,46_=1.119, *p*=0.2956; time: F_2.67,122.9_=7.03, *p*=0.0013), raw premature responses (Figure 2C_5_, HPvLP: F_1,46_=0.7796, *p*=0.3818; time: F_3.41,157.0_=4.471, *p*=0.0032), or percentage of premature responses (Figure S3I, HPvLP: F_1,46_=0.6390, *p*=0.4282). Responding for SM was also unrelated to higher versus lower consumption (Supplementary Figures 2 & 3). Importantly, these findings suggest that HP and LP mice had similar ability to learn and perform for a high-value reward, and thus that reduced alcohol responses in LP mice did not reflect differences in basic reward behavior, but, instead, a difference in engagement in responding for alcohol.

### 3.4 Reward latency is increased during alcohol sessions, but not in SM, due to occasional longer reward latency trials

Reward latency, the time from giving a correct response to entering the reward tray, is a critical metric thought to identify motivation for the reward, with faster latency taken to indicate higher drive (39). However, our initial analyses found that reward latency was not different between HP and LP mice during early-stage (Figure 3A, HPvLP: F_1,46_=0.9441, *p*=0.3363) or late-stage (Figure 3B, HPvLP: F_1,46_=12.00, *p*=0.2790) sessions for alcohol, or during strawberry milk sessions (Figure 3C, HPvLP: F_1,46_=0.0173, *p*=0.8959). Furthermore, when averaging the reward latency of the final five ethanol sessions against the five SM sessions, reward latencies were significantly slower for alcohol compared with strawberry milk: HP mice had longer latencies for alcohol compared to HP responding for strawberry milk (HP-SM, Figure 3D, t[46]=3.706, *p*=0.0006). Similarly, LP mice had longer latencies for alcohol compared to LP-SM (Figure 3D, t[45]=3.719, *p*=0.0006). Also, HP and LP mice had similar reward latencies during SM testing (Figure 3D, t[46]=0.3285, *p*= 0.7440).

**Figure 3.**
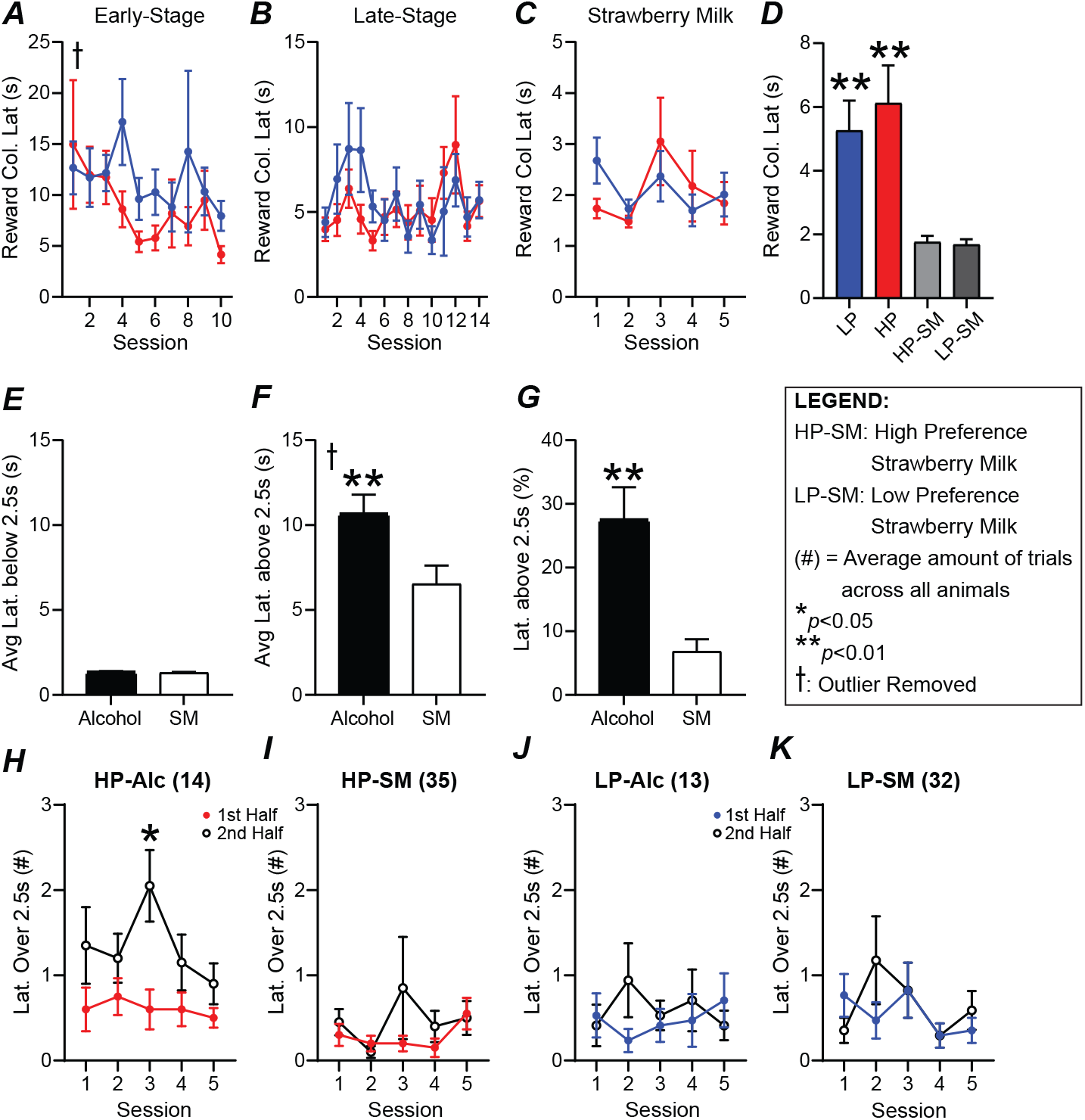
Reward latencies are longer during the second half of HP sessions for alcohol, not SM. No differences in (A) early-stage, (B) late-stage, or (C) SM reward latency. (D) Average reward latency was greater for alcohol compared to (HP/LP) SM sessions (HP-SM/LP-SM). (E) Average latencies below 2.5s were similar for alcohol and SM. (F) Average latency above 2.5s were significantly greater during alcohol sessions. (G) The percentage of latencies greater than 2.5s were much higher during alcohol sessions. HP reward latencies over 2.5s occurred significantly more during the second half of (H) alcohol, but not (I) SM sessions. LP reward latencies were similar between the first and second half during (J) alcohol and (K) SM sessions. n=48 for A-G, n=17-21/group H-K. \**p*<0.05, \*\**p*<0.01 for group effect. All data are expressed as ±standard error mean. ƚ = removed one outlier (over 200s).

To better understand potential differences in reward latency, we examined the distribution of such latencies. Indeed, we found that alcohol reward latencies could be separated into several time intervals, those more similar to SM, and others that were much longer. Specifically, when comparing average latencies under 2.5s, latency length was similar and quick when responding for alcohol or SM (Figure 3E, t[90]=0.6917, *p*=0.4909). In contrast, the average length of reward latency above 2.5s was significantly greater during alcohol testing (Figure 3F, t[90]=2.666, *p*=0.0091). There were also significantly more longer-latency (>2.5s) responses during alcohol versus SM sessions (Figure 3G, t[94]=3.794, *p*=0.0003). To better understand these longer latencies across a session, we identified whether they occurred in the first half or second half of a session (relative to number of completed trials). HP mice had significantly more longer latencies during the second half of the session for alcohol (Figure 3H, 1^st^-v-2^nd^-half: F_1,38_=6.318, *p*=0.0163), while LP mice did not (Figure 3I, 1^st^-v-2^nd^-half: F_1,32_=0.1731, *p*=0.6801). In addition, longer-latency responses for SM were fewer than for alcohol for HP and were equally distributed throughout the session in HP mice (Figure 3J, 1^st^-v-2^nd^-half: F_1,38_=1.332, *p*=0.2556) and LP mice (Figure 3K, 1^st^-v-2^nd^-half: F_1,32_=0.243, *p*=0.089). Cumulatively, these data suggest that HP mice exhibited a decrease in motivation for alcohol in the second half of the session, perhaps where mice getting more alcohol within the 5-CSRTT task participate less later in the session (addressed further in Section 3.5 and Supplementary Discussion).

When reward latencies for alcohol were examined by consumption, HC and LC mice were not different during early-stage (Figure S2A_5_, HCvLC: F_1,46_=0.0213, *p*=0.8846), late-stage (Figure S2B_5_, HCvLC: F_1,46_=0.1225, *p*=0.7280; time: F_6.30,289.3_=2.159, *p*=0.0441), or during strawberry milk sessions (Figure S2C_5_, HCvLC: F_1,46_=0.1658, *p*=0.6858). Since sorting the mice by preference proved to be more sensitive toward overall performance, consumption analysis were largely discontinued at this point.

### 3.5 Responses for alcohol, not SM, in HP mice occur more within the first half of a session

As noted in Section 3.4, changes in responding across a session may indicate altered drive for reward, e.g. where time-related shifts in reward latency in Figure 3 might relate to satiety, and such motivational changes across a session could be expressed in other measures such as less responding and/or less accurate responding. Representative trial-by-trial sessions visually describe clear differences in performance based on preference and reward (alcohol or SM, Figure S7). Thus, we investigated differences in correct, incorrect, and omissions in the first versus second half of each session to confirm any behavioral shifts. Mice that averaged at least 10 correct responses across the last five alcohol late-stage training sessions were included in this analysis, in order to more clearly assess time-related changes in performance for alcohol or SM.

HP mice displayed significantly more correct responses in the first half versus second half of sessions (Figure S6A_1_, 1^st^-v-2^nd^-half: F_1,38_=9.025, *p*=0.0047) whereas LP mice had similar correct responses in both halves (Figure S6A_2_, 1^st^-v-2^nd^-half: F_1,32_=0.6845, *p*=0.4142). In contrast, the number of incorrect responses were similar between the first and second halves for both HP mice (Figure S6C_1_, 1^st^-v-2^nd^-half: F_1,38_=0.0008, *p*=0.9773) and LP mice (Figure S6C_2_, 1^st^-v-2^nd^-half: F_1,32_=0.6345, *p*=0.4316; time: F_3.56,114_=4.254, *p*=0.0043). In addition, while there were fewer overall omissions by later training, both HP (Figure S6E_1_, 1^st^-v-2^nd^-half: F_1,38_=23.08, *p*<0.0001) and LP (Figure S6E_2_, 1^st^-v-2^nd^-half: F_1,32_=4.482, *p*=0.0421) mice displayed higher omissions in the second half then the first half. In contrast to alcohol, during SM sessions there were no differences between first and second halves in correct responding in HP (Figure S6B_1_, 1^st^-v-2^nd^-half: F_1,38_=1.232, *p*=0.2739; time: F_3.34,126.9_=8.077, *p*<0.0001) or LP mice (Figure S6B_2_, 1^st^-v-2^nd^-half: F_1,32_=0.1309, *p*=0.7199; time: F_2.35,75.24_=19.42, *p*<0.0001), or in incorrect SM responses in HP (Figure S6D_1_, 1^st^-v-2^nd^-half: F_1,38_=2.895, *p*=0.0970; time: F_3.38,128.6_=6.412, *p*=0.0002) or LP mice (Figure S6D_2_, 1^st^-v-2^nd^-half: F_1,32_=2.195, *p*=0.1483; time: F_2.98,95.44_=12.96, *p*<0.0001). However, HP mice displayed more omissions in the second half (Figure S6F_1_, 1^st^-v-2^nd^-half: F_1,38_=5.703, *p*=0.0220; time: F_2.833,107.7_=4.249, *p*=0.0081) and LP mice did not (Figure S6F_2_, 1^st^-v-2^nd^-half: F_1,32_=2.195, *p*=0.1483; time: F_2.803,89.71_=8.125, *p*=0.0001; interaction: F_4,128_=2.507, *p*=0.0453); however, the omissions difference in HP mice when responding for alcohol was p<0.0001, while the comparable difference for SM was *p*=0.0220. Together, these findings concur with reward latency results (Figure 3) that alcohol engagement in HP mice was greater during the first half of the session, which was overall not seen in LP mice or for SM responding in HP or LP, and we speculate that the second half decline in HP performance could reflect intoxicating effects of alcohol, satiety, or other factors (see Supplementary Discussion).

### 3.6 Chronic alcohol exposure enhances behavioral engagement especially in previously low-engagement individuals

For Phase 2 of our studies, mice were allowed to drink alcohol under an Intermittent Access two-bottle choice (IA2BC) drinking paradigm, with 24-hr access to 20% alcohol (versus water), three times a week, for three weeks. We were particularly interested in the possibility that IA2BC would not only enhance overall performance for alcohol, but specifically increase performance of LP mice. This could indicate that excessive consumption is particularly hazardous for individuals who innately have lower engagement with alcohol (while higher-engagement individuals already have greater risk for developing problem drinking).

Overall, HP and LP mice (defined by their alcohol behavior in initial DID sessions) had similar IA2BC consumption (Figure S4O, HPvLP: F_1,46_=0.8358, *p*=0.3654; time: F_3.93,178.3_=37.81, *p*<0.0001) and preference (Figure S4N, HPvLP: F_1,46_=3.373, *p*=0.0728; time F_7.81,355.8_=11.25, *p*<0.0001). However, HP did have greater preference during the first five sessions of IA2BC (Figure S4N, HPvLP: F_1,46_=4.479, *p*=0.0398; time F_3.24,149.1_=20.91, *p*<0.0001), although consumption levels did not differ (Figure S4O, HPvLP: F_1,46_=1.417, *p*=0.2401; time F_1.89,87.01_=49.28, *p*<0.0001).

To better assess how IA2BC drinking might influence responding in HP and LP mice, we averaged measures in the last 3 late-stage alcohol sessions and compared them with the average of the last 3 post-IA2BC sessions. While pre-IA2BC versus post-IA2BC correct responses were not different for HP (Figure 4A_1_, *p*=0.0995), IA2BC experience greatly increased correct responses in LP mice (Figure 4A_2_, *p*<0.0001). In addition, the change in correct responses (pre versus post IA2BC) was significantly different between LP and HP mice (Figure 4A_3_, *U*=176, *p*=0.0202). In contrast, HP mice had lower correct responses during SM sessions (Figure 4B_1_, *p*=0.0208), while LP mice did not (Figure 4B_2_, *p*=0.2367), nor were there differences in relative change in correct (Figure 4B_3_, *p*=0.4170). Thus, after IA2BC, LP showed significantly greater alcohol engagement, as indexed by number of correct responses, while HP mice showed less impact of IA2BC experience in this measure.

**Figure 4.**
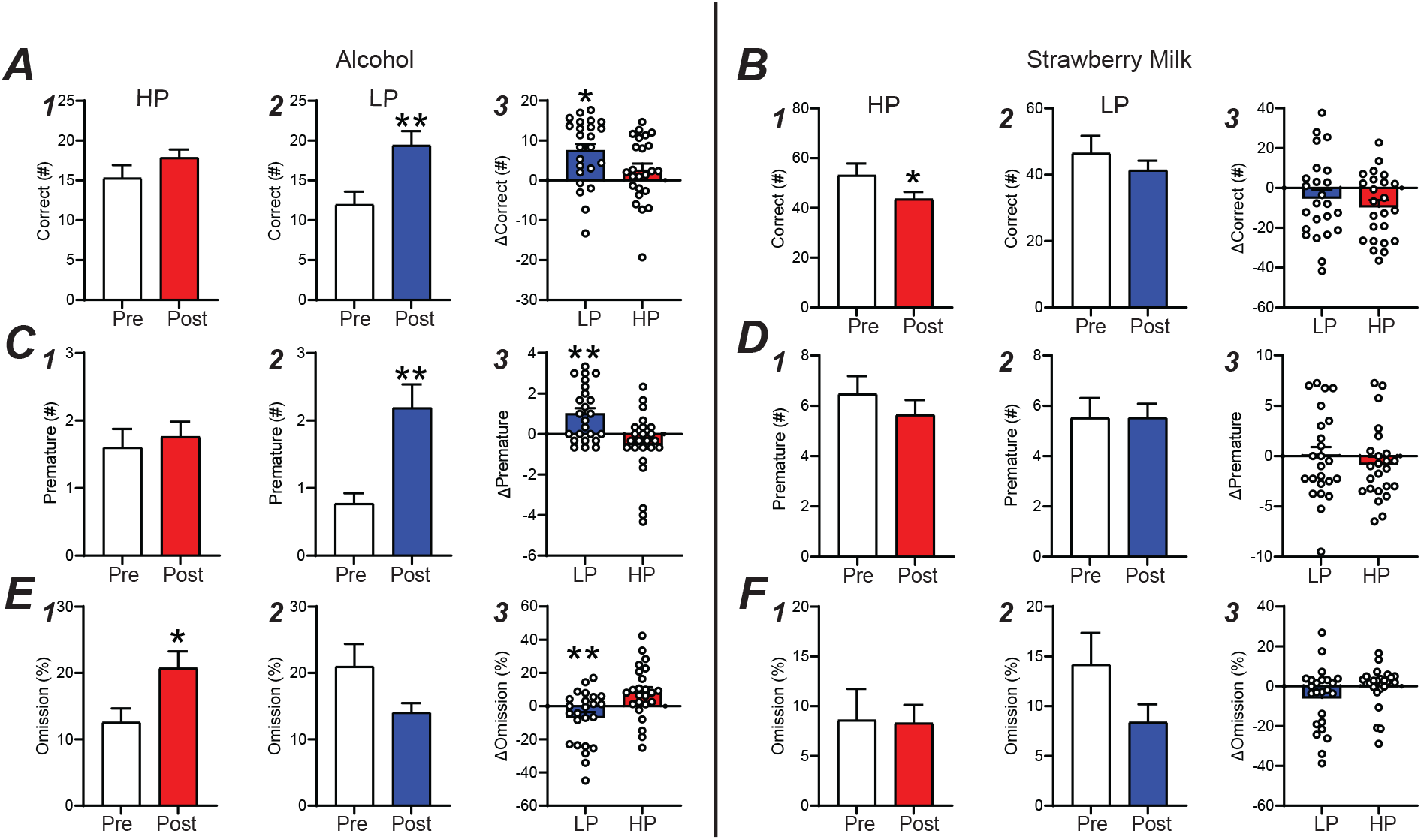
Chronic alcohol exposure promotes responding in the 5-CSRTT. (A) LP mice gave more correct responses after IA2BC and had an overall greater change in correct than HP mice. (B) HP mice displayed a decrease in correct and had a similar change in performance as LP mice during SM sessions. (C) LP premature responding greatly increased post-IA2BC and had a greater performance change than HP mice, this was not seen during (D) SM sessions. (E) %Omissions were greater in HP post-IA2BC and LP mice had a greater performance change than HP mice. (F) No differences in %omission was found during SM sessions. n=24/group, \**p*<0.05, \*\**p*<0.01 for effect of group. All data are expressed as ±standard error mean.

Similar to correct, premature responses for alcohol were unaffected by IA2BC in the HP (Figure 4C_1_, *p*=0.6017) but were increased in LP (Figure 4C_2_, *p*=0.0002) mice, with LP mice having significantly greater premature responding after IA2BC compared to HP (Figure 4C_3_, LP *U*=141.5, *p*=0.0019). SM premature were unaffected by IA2BC in HP (Figure 4D_1_, *p*=0.2832) and LP (Figure 4D_2_, *p*=0.9908), and with no differences in the change in performance after IA2BC between HP and LP (Figure 4D_3_, t[46]=0.7155, *p*=0.4779). While premature responding can indicate increased impulsivity, we speculate that increased number of premature actions may occur in parallel with overall greater responding in the task, i.e. that premature responses may also reflect the level of behavioral engagement (and that increased premature in LP mice after IA2BC emphasizes their newfound readiness to respond for alcohol compared to pre-IA2BC, see Discussion).

Finally, IA2BC did increase omissions for HP (Figure 4E_1_, *p*=0.0126), with the opposite trend for LP (Figure 4E_2_, *p*=0.1434), and LP had less loss of engagement for alcohol relative to HP (Figure 4E_3_, t[46]=3.24, *p*=0.0022), indexed by these omissions. No changes were found in HP (Figure 4F_1_, *p*=0.2076) or LP (Figure 4F_2_, *p*=0.2182) during SM sessions, and relative performance changes were also similar between the groups (Figure 4F_3_, *U*=208, *p*=0.1015). Additional measures are detailed in Supplementary Results and Figure S8.

Together, these data strongly suggest that IA2BC overall increased engagement with alcohol responding significantly more in LP versus HP mice. Interestingly, in LP mice, IA2BC promoted responding even in mice who, pre-IA2BC, had barely responded (5 or fewer correct pre-IA2BC, increasing to ∼15 correct post-IA2BC). IA2BC also had limited impact on SM sessions, with only HP mice displaying decreased correct SM responses (although no decrease in alcohol responses). However, this SM decrease may also suggest decreased sucrose seeking behaviors after chronic alcohol exposure as seen in previous studies (27). The effect of chronic alcohol exposure on performance, regardless of preference, opens many avenues of investigation.

## 4 Discussion

Excessive alcohol consumption is a widely prevalent activity that may promote the development of AUD, where alcohol intake becomes a necessity for the individual and becomes a considerable barrier to treatment. Increased motivation and abnormal attentiveness for alcohol are often signs of AUD, and individuals with higher trait impulsivity are also at greater risk for AUD (see Introduction). Thus, it is critical, when seeking to develop novel treatments, to discover biological mechanisms that promote these forms of behavioral engagement for alcohol. In the current study, we used the 5-CSRTT, a behavioral paradigm that can measure a number of facets of behavioral engagement (e.g. attention, impulsivity, and motivation) in the same session. Importantly, we, for the first time, adopted the 5-CSRTT to train mice to respond for alcohol as reward. All previous studies examined how alcohol exposure alters 5-CSRTT responding for sugar, but we wanted the mice to associate the task with alcohol, rather than sugar, so that future implementation of more challenging forms of the task (e.g. to assess impulsivity under variable timing), will reflect their motivation and overall performance for alcohol. This removes the risk of habit formation in responding for sugar, which could impact responding for alcohol if the reward was switched from sugar during training to alcohol during testing. Interestingly, using this novel alcohol 5-CSRTT paradigm, we found that alcohol preference was a more sensitive indicator of performance for alcohol in the 5-CSRTT, rather than consumption. HP mice learned the task faster and had greater participation than LP mice. Further differences were found within sessions, where HP mice had more correct responses and faster latencies in the first half of the session, perhaps suggesting a form of “front-loading” behavior and/or satiety later in the session (see below). Finally, we found that three weeks of alcohol drinking (IA2BC) greatly promoted responding for alcohol in LP mice, with lesser or no change in HP related to IA2BC. This suggests that IA2BC produced a greater increase in motivation for alcohol in mice that innately began with lower preference, while innately higher preference mice were already more alcohol-responsive. Importantly, IA2BC overall had little effect on SM sessions, further underscoring the specificity in IA2BC effects on increasing alcohol engagement in LP individuals.

Studies have used the 5-CSRTT to compare treatment effects on motivation, attention, and impulsivity for sugar (40, 41). Indeed, alcohol vapor, liquid diet, and gavage treatment have all been used to induce increase premature responding or attentional errors in the 5-CSRTT when responding for sugar (25, 27, 30, 42). However, to date there are no studies we are aware of that exclusively use an intoxicant as reward for 5-CSRTT. One goal of this study was to effectively train mice in this complex task for an alcohol-only reward (10% alcohol), with no other additives (such as saccharin or even saccharin fade). This important advancement has allowed us to observe how motivated an animal was to wait during the intertrial interval, give a correct response, then retrieve the alcohol. Previous studies have trained animals to respond for alcohol in other behaviors such as differential reinforcement of low rates of responding (DRL) and progressive ratio tasks (43, 44). While progressive ratio is valuable for measuring motivation, and DRL for impulsivity, the 5-CSRTT is designed to assess a broader range of measures in the same session. However, training animals in the 5-CSRTT with alcohol remains largely unexplored. We detail the nuances that come with having an intoxicant as a behavioral reward in this task in the Supplemental Discussion, potential effects of increasing intoxication. Thus, we suggest that 5-CSRTT with alcohol as the reward can be robustly studied. This method of alcohol responding will likely be invaluable for future studies of how different interacting factors lead to different pathways to excessive drinking, including conditions with higher challenge (e.g. requiring greater attention or waiting) that some variants of 5-CSRTT testing can examine.

We find that 5-CSRTT performance was significantly related to alcohol preference rather than consumption, including where HP mice demonstrated significantly more engagement with alcohol compared with LP mice. One speculation is that preference is related to an innate attention to and engagement with some condition. In this model, HP mice may reflect an individual who repeatedly orders a drink when there is an inherent waiting period (bartender order to delivery of alcohol), payment (correct response), and, finally, reward retrieval (consumption of drink). LP mice, however, may be compared with a more social drinker, where they drink when alcohol is easily available (2-bottle choice), but are more likely to be dissuaded if it requires actively work for it (purchasing, waiting, or traveling for alcohol). To further speculate, it is interesting that, in addition to high trait impulsivity, people at risk for binging have higher alcohol preference (8-10). However, determining clear relations in human studies remains challenging, and the ability to control factors in rodent studies has given insights into the alcohol-impulsivity relationship. For example, mice genetically selected for high alcohol preference display higher impulsivity in a delay discounting task compared with mice selected for low preference, although some aspects of impulsivity (amphetamine and lithium reduction of impulsivity) are not related to preference (45, 46). In addition, mice genetically selected for high alcohol consumption displayed impaired response inhibition in a Go/No-go task but were not different from low consumption in delay discounting (47). However, we should note that consumption level is still a very important factor, and several studies have assessed differential neural mechanisms that underlie higher versus lower intake level (38, 48). Thus, we emphasize that HP and LP had similar drinking levels that, based on blood alcohol assessments in our previous mouse studies (49, 50), both HP and LP drank sufficient alcohol to on average reach binge level. In addition, premature responding in the 5-CSRTT is an indicator of impulsivity, and we found that HP mice had higher premature responses during early-stage training. This may suggest that their propensity for alcohol promotes error in the ability to wait (for the stimulus), although we speculate that greater premature responses along with greater overall responding could in some cases reflect greater engagement rather than impulsivity per se. Future studies will be needed to further dissect these and other aspects of behavior that promote excessive drinking. Cumulatively, the field continues to make substantial strides toward understanding the relationship between impulsivity, alcohol preference and intake, but the interconnection of these remains ambiguous.

Impulsivity is a complex construct that has come under scrutiny due to the wide breadth of behaviors it encompasses (12-14). Risk-taking tasks, delay-discounting, DRL, reversal learning, Go/No-Go, and, here, the 5-CSRTT are examples of impulsivity-related tasks, however they all measure impulsivity in different ways. Impulsive behaviors have traditionally been separated into two domains, impulsive action and Impulsive choice (51-54). Under impulsive action, Go/No-Go and reversal learning measure action inhibition while the 5-CSRTT measures the ability to wait. In contrast, impulsive choice tasks measure sensitivity to delayed (e.g. delay-discounting) and risky choices (e.g. Balloon analogue risk task). We have chosen the 5-CSRTT since the task allows assessment of premature responses (related to impulsivity) as well as omissions and accuracy (more clear indicators of engagement). However, premature responses during standard training sessions may describe participation opposed to impulsivity. Future studies will utilize randomized waiting periods to truly test impulsive responding. Thus, with 5-CSRTT we have the ability to measure impulsivity (among other measures) related to voluntary alcohol acquisition, especially in future (and ongoing) studies using 5-CSRTT variants that more clearly assess (e.g. variable timing of reward presentation). Importantly, even with the broad nature of impulsivity, clinical studies using impulsivity tasks often shown its relationship to alcohol use (33-37) and the 5-CSRTT has been used clinically to predict higher alcohol intake in highly impulsive individuals (26). Here, we attempted to bridge a much-needed gap in rodent alcohol-impulsivity studies so that future work can focus on impulsive action for voluntary alcohol consumption, and related behavioral indicators of excessive intake, to decipher potential patterns and biomarkers that mirror clinical findings.

Several studies have shown how disruptions in various cortical areas can alter behavioral engagement detected through the 5-CSRTT (28, 55, 56). The anterior cingulate cortex (ACC), a region involved in higher-level cognitive function, modulates accuracy and omissions (55, 57) and is required for top-down action control within the 5-CSRTT (58). Further, errors in decision-making after chronic alcohol use have been linked to ACC (59-61). Additionally, the anterior insula (aINS) has been implicated in impulsive behaviors (62-64), including where premature responding in the 5-CSRTT correlates with aINS thinning in rat (65). The aINS has also been heavily associated with attention (66-69), as a critical part of the salience network, and been implicated in problem alcohol drinking (70-74). Though there are many regions associated with behavioral engagement, the ACC and aINS may be future targets of investigation linking chronic alcohol intake and changes in engagement for alcohol.

Currently, investigators may be hesitant to replace the typical appetitive reward with an intoxicant. However, in the present study, we observed the satisfactory performance of mice while training in the 5-CSRTT for an alcohol reward. This paradigm displays the relationship between motivation for freely accessible alcohol (DID, 2-bottle choice) and for when mice must perform an attention-based task (5-CSRTT) for alcohol. Interestingly, HP mice were more likely to perform the task than their LP counterparts, while protracted alcohol consumption increased engagement with alcohol in LP mice significantly more than HP, while SM responding was higher than for alcohol and overall not different in HP versus LP or before and after IA2BC. In addition, differences in performance found within HP sessions emphasize the importance of doing a trial-by-trial analysis to maximize the efficacy and interpretability of the data. Though we do not have a non-IA2BC control, the differences between HP and LP mice after IA2BC found within this study detail the risk chronic alcohol intake has on problem drinking even in those that did not prefer alcohol. Thus, our behaviorally-focused series of experiments set a foundation to answer important neurological hypotheses that will be featured in future studies. Cumulatively, our findings offer a novel insight into preference-related motivation for and engagement with alcohol, that can be used to identify behavioral patterns and brain mechanisms among the different factors (including attention, reward motivation, impulsivity, and perseveration) that could come together in different ways to promote excessive alcohol drinking and its substantial harms.

## Abbreviations

5-CSRTT: 5-choice serial reaction time task
HP: high preference
LP: low preference

## ACKNOWLEDGEMENTS

This work was funded by T32 AA 007462 “Training Grant on Genetic Aspects of Alcoholism”. We would also like to thank Raziel Fraser for editing the paper and Desarae Dempsey for analysis insights.

## AUTHOR CONTRIBUTION

PS, DM, HM designed the behavioral schedule and collected the data. PS, DM analyzed the data. PS and FH wrote the manuscript. All authors provided critical review of the content and approved of the final version for publication.

