## Supplementary Material for "Low alcohol preferring mice have reduced task engagement during a waiting task for alcohol, which is enhanced by intermittent alcohol drinking"

**SUPPL. METHODS**

**2.2 5-Choice serial reaction time task (5-CSRTT)**

**(A) Water Restriction and Apparatus.** Mice were placed on water restriction throughout the experiment to promote performance for the alcohol reward (1). Mice were given two hours of water access after their behavioral sessions. All operant training procedures were performed in a noise-attenuating Bussey-Sakisda Rodent Touch Screen Chamber (Lafayette Instruments Co., Lafayette, IN) designed for mice. Briefly, the chamber has a perforated stainless-steel floor with trapezoidal plastic walls enclosed with a touchscreen (12.1 in, resolution 800 x 600) with a plastic mask outlining 5 4 x 4 cm holes spaced 1 cm apart and 1.5 cm from the floor. Infrared photo-detection beams are used to detect responses to the touchscreen and for entrances to the rear reward collection area. Additionally, the chambers are equipped with a 3W light for punishing responses, a speaker to deliver 65 dB tone for correct responses, and a ventilation fan. Mice could be observed through an overhead infrared-sensitive camera. An illustration can be seen in figure 1B.

**(B) Pre-training.** (Figure 1A) Mice were first habituated to the operant chambers for 15 min for two days. During these sessions, the chambers deliver 20ul of reward after each tray exit to promote familiarity with the reward location. The next three days consist of a Fixed-Ratio 1 (FR1) schedule for a 10% alcohol reward. These sessions lasted 60 min and had an unlimited number of trials. Mice performing earlier in the day showed decreased performance compared to the mice being tested later in the day, thus we added an additional FR1 session reversing the order the animals were tested. There was no change in performance in any of the animals.

**(C) Early-Stage Training.** (Figure 1A) Mice were then transitioned into performing an easy version of the 5-CSRTT. Mice are placed into a dark chamber and the program begins with 200ul of free reward (10% alcohol) illuminated in the reward trough in the rear of the arena. Each trial is initiated with a head entrance into the reward tray. Once initiated, the 5s intertrial interval (ITI) begins. Once the ITI ends, one of the 5 square holes will illuminate for 10s (stimulus duration, SD). The animal has an additional 37s to give a response (limited hold, LH) before the trial is considered an omission. The animal can make several different responses. A touch on one of the five holes during the ITI period will result with a punishment (flash of light), recorded as a premature response, and will reset the ITI timer. If the stimulus is presented and the animal touches one of the unlit holes, there will be a punishment and it will be recorded as an incorrect response. After an incorrect or correct response is given, any additional responses in the five holes are recorded as a perseverative response. If a correct response is given, 20ul of 10% alcohol will dispense from a liquid pump in the illuminated reward tray (Figure 1B). After a correct, incorrect, or omission response, the reward tray will illuminate until the next tray exit, thereby initiating the next trial. Mice continued this training for 10 days. A flow chart outlining the pathways of this task can be seen in supplemental figure 1.

**(D) Late-Stage Training.** (Figure 1A) As mice continued training, performance of both groups began to merge and classified this as late-stage training. Mice spent 14 days training in the 10s SD before being taken off water restriction to observe change in performance. After one week with *ad libitum* water access, the mice were weight restricted to be tested for one week at 10s SD and 5s ITI for a strawberry milk (Nesquik, Nestle) reward to compare performance between the 10% alcohol and a traditional 5-CSRTT reward.

**2.3 Two-Bottle Choice and Intermittent Access**

**(A)** Throughout 5-CSRTT training, mice were given weekend access to 10% alcohol for 2 h to promote response to the reward (Figure 1A, Phase 1). Custom-built, low drip sipper tubes were used to reduce dripping from overactive mice that may climb on the cage. These tubes consisted of a Falcon 15 mL conical tube (Fisher Scientific, Hampton, NH) with the bottom cone cut and filed down. A sipper (Ancare Corp, Bellmore, NY) was then placed inside the tube and shrink-wrapped using a heat gun. A rubber stopper (size:0#, StonyLab, Nesconset, NY) was used to plug the opposite end. Bottles were weighed before and after sessions and consumption was calculated using the weekly weight of the mouse. Preference was calculated as the total amount of alcohol consumed divided by total liquid consumed.

**(B)** Similar to Lei et al. 2019 (2), mice were given 24 h access to 20% alcohol every Sunday, Tuesday, and Thursday starting at 7:00 am (Figure 1A, Phase 2). Consumption was calculated using the weekly weight of the mouse. During non-alcohol days, two-bottles filled with water were present to maintain familiarity with the bottles.

**(C)** Post-Intermittent Access Testing. (Figure 1A, Phase 2) Mice were given 4 sessions for alcohol reward and 4 for a SM reward using a 10s SD and 5s ITI duration. Importantly, mice continued intermittent access throughout the testing, resulting in approximately six weeks of total intermittent access. Mice were tested between 5-9 hours after their last alcohol access session to ensure the mice were performing during adequate acute withdrawal. Water bottles were not given until after the behavior was completed for that animal on that day. For analysis we excluded the first day to remove potential burst in behavior from reintroducing the mice to the task.

**SUPPL. RESULTS**

**3.1. Mice with high alcohol preference or consumption show higher engagement in early-stage training.**

Alcohol preference and consumption were correlated across all mice (*p*<0.0001, R^2^=0.4566, Figure S1B). A correlation was not found between preference and consumption of HP mice (Figure S1C, *p*=0.4776, R^2^=0.0232), but a correlation was found in LP mice (*p*=0.0002, R^2^=0.4859). A correlation between preference and consumption was found in LC mice (Figure S1D, *p*<0.0001, R^2^=0.473) but not for HC mice (*p*=0.2018, R^2^=0.089).

HC and LC raw premature responses were similar between both groups (Figure S3A, HCvLC: F_1,46_=2.68, *p*=0.1085; time: F_1.36,62.56_=4.11, *p*=0.0350; interaction: F_9,414_=2.19, *p*=0.0219) with no post-hoc significance. The percentage of premature responses was also similar between the groups (Figure S3B, HCvLC: F_1,46_=2.042, *p*=0.1598; time: F_2.54,116.6_=2.90, *p*=0.0465; interaction: F_9,414_ = 2.70, *p*=0.0046) with no post-hoc significance. This data may not suggest traditional impulsive responding but greater participation in learning the task.

**3.2 Preference predicts performance, while alcohol consumption does not, during late-stage training.**

In HC and LC SM responding, all had similar number of trials (Figure S2C_1_, HCvLC: F_1,46_=0.69587, p=0.4085; time; F_2.78,128_=42.59, *p*<0.0001; interaction: F_4,184_=3.167, *p*=0.0151), accuracy (Figure S2C_2_, HCvLC: F_1,46_=0.0346, *p*=0.8532; time: F_2.87,132_=7.478, *p*=0.0001), correct responses (Figure S2C_3_, HCvLC: F_1,46_=0.1998, *p*=0.6570; time F_2.75,126.5_=44.69, *p*<0.0001; interaction: F_4,184_=3.101, *p*=0.0168), omissions (Figure S2C_4_, HCvLC: F_1,46_=0.1219, *p*=0.7286; time: F_2.69,123.8_=6.968, *p*=0.0004), raw premature responses (Figure S3C, HCvLC: F_1,46_=0.0145, *p*=0.9046; time: F_3.461,159.2_=4.651, *p*=0.0024), or percentage of premature responses (Figure S3D, HCvLC: F_1,46_=0.0793, *p*=0.7795; time: F_3.29,150.5_=1.538, *p*=0.2032; interaction: F_4,183_=2.644, *p*=0.0351).

The first day of training following a two-day alcohol session displayed a burst of behavior that decreases the following days. We opted to see if omission of these burst days affected the overall result. HP mice displayed increases in trials (Figure S4A, HPvLP: F_1,46_=6.454, *p*=0.0145; time: F_6.87,316.3_=2.229, *p*=0.0327). Accuracy was similar between the groups (Figure S4B, HPvLP: F_1,46_=1.669, *p*=0.2028; time: F_6.39,293.9_=5.416, *p*<0.0001). The number of correct responses was higher in HP mice (Figure S4C, HPvLP: F_1,46_=4.946 *p*=0.0311; time: F_6.54,301.0_=3.376, *p*=0.0023; interaction: F_10,460_=2.215, *p*=0.0160), with post-hoc significance on the second session. Omissions also remained similar between HP and LP mice (Figure S4D, HPvLP: F_1,46_=1.942, *p*=0.1701, time: F_5.85,269.2_=3.358, *p*=0.0036). Raw premature responses were similar between both groups (Figure S4E, HPvLP: F_1,46_=2.943, *p*=0.0928). The percentage of premature responses was also similar between the groups (Figure S4F, HPvLP: F_1,46_=2.877, *p*=0.0966). When analyzing by consumption, HC and LC mice had similar number of trials (Figure S4G, HCvLC: F_1,46_=1.935, *p*=0.1709; time: F_6.83,314.2_=2.229, *p*=0.0331). Accuracy was similar between the groups (Figure S4H, HCvLC: F_1,46_=0.0797, *p*=0.7790; time: F_6.4,294.5_=5.354, *p*<0.0001). The number of correct responses was also similar between groups (Figure S4I, HCvLC: F_1,46_=0.8195 *p*=0.3701; time: F_6.48,298.2_=3.333, *p*=0.0026). Omissions also remained similar between HP and LP mice (Figure S4J, HCvLC: F_1,46_=0.0935, *p*=0.7611; time: F_5.78,266.0_=3.383, *p*=0.0035). Raw premature responses were similar between both groups (Figure S4K, HCvLC: F_1,46_=0.7176, *p*=0.4013). The percentage of premature responses was also similar between the groups (Figure S4L, HCvLC: F_1,46_=0.6574, *p*=0.4217; interaction: F_10,460_=1.904, *p*=0.0426). In concurrence with the full data set, where the “burst” days are included, these sessions of increased behavior do not change the overall effect.

There were four mice that had alcohol preference which rounded to 79% but were split into both HP and LP groups. Thus, we removed these to see if it would impact our results. HP mice still displayed increases in trials (Figure S3J, HPvLP: F_1,42_=4.963, *p*=0.0313; time: F_7.49,3134.4_=17.53, *p*<0.0001). The number of correct responses was higher in HP mice (Figure S3K, HPvLP: F_1,42_=4.618, *p*=0.0375; time: F_7.379,309.9_=7.741, *p*<0.0001). Accuracy was similar between the groups (Figure S3L, HPvLP: F_1,42_=2.737, *p*=0.1055; time: F_7.42,311.7_=5.996, *p*<0.0001). Omissions also remained similar between HP and LP mice (Figure S3M, HPvLP: F_1,42_=3.057, *p*=0.0877; time: F_6.37,267.6_=2.976, *p*=0.0067). Since these four mice did not drastically impact our results in late-stage training, we opted to include them in our experiments.

Lastly, mice were given one week of training while given unlimited water access to observe the impact of water restriction on performance of HP and LP mice. While on water access, HP mice performed a greater number of trials than LP mice (Figure S5A, HPvLP: F_1,46_=4.055, *p*=0.0499). Both groups had similar accuracy (Figure S5B, HPvLP: F_1,46_=1.926, *p*=0.1719). HP mice displayed a greater number of correct responses (Figure S5C, HPvLP: F_1,46_=5.690, *p*=0.0212; time: F_2.93,134.6_=3.355, *p*=0.0218), but had similar percentage of omissions as LP mice (Figure S5D, HPvLP: F_1,46_=2.331, *p*=0.1337). Reward latency was also similar between HP and LP mice (Figure S5E, HPvLP: F_1,46_=1.291, *p*=0.7210). When analyzed by consumption, HC and LC mice had similar number of trials (Figure S5F, HPvLP: F_1,46_=2.172, *p*=0.1473), accuracy (Figure S5G, HPvLP: F_1,46_=0.0064, *p*=0.9365), correct (Figure S5H, HPvLP: F_1,46_=0.8620, *p*=0.3580; time: F_2.92,134.5_=3.421, *p*=0.0201), omissions (Figure S5I, HPvLP: F_1,46_=0.0132, *p*=0.9089) and reward latency (Figure S5J, HPvLP: F_1,46_=0.3739, *p*=0.5439). These results suggest that water access did not change the overall effects found between the groups.

**3.6 Chronic alcohol exposure enhances behavioral engagement.**

The number of correct responses was not different post-IA for HP (Figure 4A1, Pre-IA: Median (Med)=16.83; Post-IA: Med=18.33, *p*=0.0995) but LP mice (Figure 4A2, Pre-IA: Mn=11.97, Std=7.90; Post-IA: Mn=19.42, Std=8.70, *p*=0.0002) greatly increased correct responses. LP (Med=9.67) mice had a greater change in correct responses than HP (Med=2.167) mice after IA (Figure 4A3, U=176, *p*=0.0202). For SM testing, the number of correct responses was also lower post-IA2BC for HP (Figure 4B_1_, Pre-IA2BC: Med=54.5; Post-IA2BC: Med: 43.0, *p*=0.0208) but not in LP mice (Figure 4B_2_, Pre-IA2BC Mn=46.47, Std=25.65; Post-IA2BC: Mn=41.36, Std=13.94, *p*=0.2367). Both groups displayed similar change in performance after IA2BC (Figure 4B_3_, t[46]=0.82, *p*=0.4170).

Omissions were greater post-IA2BC in HP mice (Figure 4C1, Pre-IA2BC: Med Med=9.15; Post-IA2BC: Med=16.32, *p*=0.0126) but was not in LP mice (Figure 4C2, Pre-IA2BC: Med=17.63; Post-IA2BC: Med=13.60, *p*=0.1434). LP mice had a greater change in omission performance after IA2BC than HP mice (Figure 4C3, t[46]=3.24, *p*=0.0022). During SM sessions, percentage of omissions was not different post IA2BC in HP mice (Figure 4D_1_, Pre-IA2BC: Med Med=54.5; Post-IA2BC: Med= 43.0, *p*=0.2076) as well as in LP mice (Figure 4D_2_, Pre-IA2BC: Med=7.28; Post-IA2BC: Med=4.71, *p*=0.2182). HP (Med=2.50) and LP (Med=-2.44) mice had similar performance changes after IA2BC (Figure 4B_3_, *U*=208, *p*=0.1015).

Premature responses were unaffected by IA in the HP (Figure 4E1, Pre-IA2BC: Med=1.13; Post-IA2BC: Med=1.75, *p*=0.6017) but were increased in LP (Figure 4E2, Pre-IA2BC: Med=0.75; Post-IA2BC: Med=1.75, *p*=0.0002) mice. LP (Med=0.83) mice had a greater change in premature responding after IA compared to HP (Figure 4E3, Med=-0.33; U=141.5, *p*=0.0019). During SM sessions, premature responses were unaffected by IA2BC in the HP (Figure 4E_1_, Pre-IA2BC: Mean (Mn)=6.47, Std=3.51; Post-IA2BC: Mn=5.65, Std=2.86, *p*=0.2832) and LP (Figure 4E_2_, Pre-IA2BC: Mn=5.52, Std=3.85; Post-IA2BC: Mn=5.53, Std=2.70, *p*=0.9908) mice, and both displayed similar changes in performance after IA2BC (Figure 4E_3_, t[46]=0.72, *p*=0.4779).

The number of trials for HP mice was greater post-IA2BC (Figure S8A_1_, Mn=29.27, Std=8.83) compared with pre-IA2BC (Mn=38.26, Std=7.22, *p=*0.0004) and also in LP mice (Figure S8A_2_, Pre-IA: Mn=23.96, Std=9.44; Post-IA: Mn=38.32, Std=10.40, *p<*0.0001). The change in performance between HP and LP after IA2BC was similar (Figure S8A_3_, t[46]=1.71, *p*=0.0949). During SM sessions, trials for HP mice were lower post-IA2BC (Figure S8B_1_, Mn=59.97, Std=13.78) compared with pre-IA2BC (Mn=68.96, Std=20.54, *p=*0.0052). No difference was observed in the LP mice (Figure S8B_2,_ Pre-IA Mn=62.21, Std=23.37; Post-IA2BC: Mn=57.75, Std=13.55, *p*=0.3676). The change in performance between HP and LP after IA2BC was similar (Figure S8B_3_, t[46]=1.12, *p*=0.2668).

Overall accuracy did not change in either group HP (Figure S8C_1_, Pre-IA2BC: Median (Med)=68.02; Post-IA2BC: Med=70.29, *p*=0.2076) or LP mice (Figure S8C_2_, Pre-IA2BC: Med=65.98; Post-IA2BC: Med=66.55, *p*=0.1280). Both groups had similar changes in performance after IA2BC (Figure S8C_3_, t[46]=0.54, *p*=0.5952). During SM sessions, accuracy did not change in either HP (Figure S8D_1_, Pre-IA2BC: Mn=74.72, Std=18.87; Post-IA2BC: Mn=74.40, Std=13.26, *p*=0.8875) or LP mice (Figure S8D_2_, Pre-IA2BC: Med=78.64; Post-IA2BC: Med: 74.86, *p*=0.6231). Both groups had similar changes in performance after IA2BC (Figure S8D_3_, HP: Med= -4.08, LP: Med=-2.21, *U*=274, *p*=0.7827).

The number of incorrect responses remained similar in HP (Figure S8E_1_, Pre-IA2BC: Mn=8.33, Std=2.25; Post-IA2BC: Mn=8.63, Std=3.01, *p*=0.6620), but were increased in LP mice (Figure S8E_2_, Pre-IA2BC: Mn=7.33, Std=2.88; Post-IA2BC: Mn=9.90, Std=2.93, *p*=0.0017). LP mice had a greater relative increase in incorrect after IA2BC than HP mice (Figure S8E_3_, t[46]=2.15, *p*=0.0367). During SM sessions, incorrect responses remained similar in HP (Figure S8F_1_, Pre-IA2BC: Mn=11.86, Std=4.50; Post-IA2BC: Mn=12.45, Std=2.89, *p*=0.3974), but were increased in LP mice (Figure S8F_2_, Pre-IA2BC: Mn=10.63, Std=3.59; Post-IA2BC: Mn=13.5, Std=3.27, *p*<0.0001), with LP mice having a greater relative increase after IA2BC than HP mice (Figure S8F_3_, t[46]=2.52, *p*=0.0152).

Percentage of premature also was similar after IA2BC in HP (Figure S8G_1_, Pre-IA2BC: Med=5.19; Post-IA2BC: Med=4.83, *p*=0.3596) but increased in LP (Figure S8G_2_, Mn=3.23, Std=2.63; Post-IA2BC: Mn=5.16, Std=4.07, *p*=0.0329) mice. LP mice had a greater change in percent premature responses than HP mice after IA2BC (Figure S8G_3_, t[46]=2.75, *p*=0.0085). During SM sessions, premature percentage for HP was similar after IA2BC (Figure S8H_1_, Pre-IA2BC: Mn=8.59, Std=3.94; Post-IA2BC: Mn=8.85, Std=4.50, *p*=0.7777) and LP (Figure S8H_2_, Mn=7.61, Std=3.80; Post-IA2BC: Mn=8.88, Std=3.66, *p*=0.2325). There were no differences in the change in performance between HP (Med=-0.31) and LP (Med=0.13) mice (Figure S8H_3_, *U*=266, *p*=0.6604).

Lastly, the total amount of tray entries increased in HP (Figure S8I_1_, Pre-IA2BC: Mn=62.25, Std=22.18; Post-IA2BC: Mn=91.79, Std=25.87, *p*<0.0001) and in LP (Figure S8I_2_, Pre-IA2BC: Mn=49.16, Std=22.9; Post-IA2BC: Mn=87.99, Std=28.79, *p*<0.0001) mice. Both groups had a similar change in tray entries after IA2BC (Figure S8I_3_, HP: Med=26.67, LP: Med=39.83, *U*=213, *p*=0.1240). During SM sessions, tray entries decreased in HP (Figure S8J_1_, Pre-IA2BC: Med=161.9; Post-IA2BC: Med=136.3, *p*=0.0009) and in LP (Figure S8J_2_, Pre-IA2BC: Med=154.1; Post-IA2BC: Med=37.92, *p*=0.0491) mice, with both groups having a similar performance change after IA2BC (Figure S8J_3_, HP: Med=-19.38, LP: Med=-26.63, *U*=262.5, *p*=0.6057).

**SUPPL. DISCUSSION**

One of our concerns while training the mice was their low accuracy. Traditionally, the 5-CSRTT criterion for sufficient performance requires greater than eighty percent accuracy and less than twenty percent omission rates (3-5). However, we have observed mice that have consistently reached fifty to sixty percent accuracy while others achieve approximately thirty. Since the reward is alcohol, we adjusted our criterion to focus more on number of trials and correct responses per session. This may be an error in attention vigilance when alcohol is the reward due to its intoxicating effects as it has been previously found that prenatal alcohol exposure and alcohol binge can dimmish attention within the 5-CSRTT (6-8). When the mice were switched to SM, they quickly escalated to the traditional criteria for this task. This data provides evidence that alcohol training was sufficient to enable the mice to understand the task but must be examined differently than sessions with a more traditional reward such as SM.

Weight restriction, where the animals are kept at 85% of the *ad libitum* weight, is used to motivate animals for the reward in many behaviors that give a caloric reward (9-13). During our experiments, we opted to use water restriction, where the animals are given only 2 hours of water each day, to motivate our mice for the alcohol which has been used in previous studies within the 5-CSRTT (1). We hypothesized that thirst was a better motivator for 10% alcohol than hunger since alcohol drinking is not a typical method for mice to receive calories. For SM testing, we switched to weight restriction as it may be the more relevant motivator due to caloric restriction. We identify that this dilutes the comparison between alcohol and SM sessions and future studies will test using weight or water restriction throughout all sessions to address this. However, the differences and performance changes in HP and LP during alcohol sessions is not affected by this factor.

During one week of late-stage training, we took the mice off water restriction to see their performance within the 5-CSRTT and for alcohol drinking in the 2-bottle choice. We found that the number of trials and correct responses decreased, however the differences found in those metrics between HP and LP remained (Figure S5). During the 2-bottle choice, we observed a drop in overall consumption (Figure 1F, session 8) with no effect on preference (Figure 1E, session 8). This data provided enough evidence that the mice should remain on water restriction throughout the experiment to maintain greater participation. Additionally, throughout the experiment our mice displayed “bursts” of behavior on Mondays, the start of each training week. Mice were given a 2-bottle choice of 10% alcohol for 2h session on the previous two days without 5-CSRTT training. The reintroduction to the 5-CSRTT each Monday may be the reason for this transient increase in responding within both groups. In Figure S4, the “burst” days were removed from the analysis to see if the effect observed was a product of this phenomenon. Indeed, the differences found in HP and LP mice in number of trials and correct responses remain while HC and LC mice remain similar throughout the training, thereby concluding that inclusion of these days were appropriate for further analysis.

The use of the Bussey-Sakisda behavioral chambers and complementary software allowed us to identify specific trial differences within a session (14-16). Mice that received an average of 100mL or more of reward in their final five sessions were used in the analyses. Low-performing mice percentages would provide an inaccurate assessment of behavior. Since the analysis is done with average percent of a variable, a low-performing animal may have extremely high or low percentages for providing few responses. As described previously, HP mice gave more correct responses and faster reward collection latencies during the first half of their sessions compared to their last half. Further, reward collection latencies, a measure of motivation for the reward, for alcohol have periodic reward latencies where the mouse delayed collection of the reward after a correct response. This data leads us to hypothesize that mice may eagerly perform for alcohol but are fighting against the effect alcohol has on them. Mice are given 200uL of alcohol at the start of the session which would increase their blood alcohol level at the start of the task. As they continue to obtain alcohol from correct responding, the mouse may hit their alcohol recovery period. This recovery period can be seen as the animal’s inherent alcohol satiety, understanding that some of the current alcohol in their system needs to be metabolized before further consumption. Studies have observed mechanisms of alcohol satiety for therapeutic avenues, here we speculate it as a possible reason for these lengthy reward latencies (17). Additionally, if the overall session averages were the only way to analyze their behavior, then we would assume the mice were spending an average of over 8 seconds after a correct response to retrieve the alcohol reward. Sucrose or milk latencies are typically less than 2.5 seconds and when comparing with the alcohol latencies an investigator might conclude the animals are not motivated for the reward (18, 19). Our trial-by-trial analysis revealed that only a small percentage of reward latencies that are over 2.5s (~30% for alcohol and ~10% for SM). When viewing the average latency below 2.5s we see that their average latency for alcohol matches SM latencies. Thus, we would like to stress the importance of trial-by-trial analysis when using an intoxicant, in this case alcohol, as the reward.

**SUPPL. FIGURE LEGENDS**

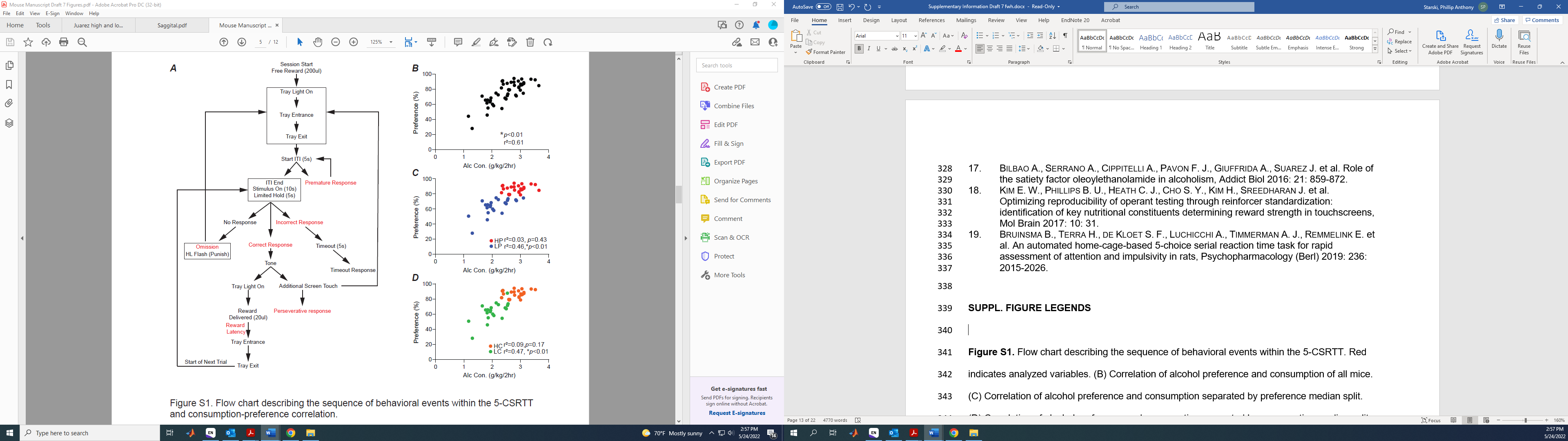

**Figure S1.** Flow chart describing the sequence of behavioral events within the 5-CSRTT. Red indicates analyzed variables. (B) Correlation of alcohol preference and consumption of all mice. (C) Correlation of alcohol preference and consumption separated by preference median split. (D) Correlation of alcohol preference and consumption separated by consumption median split. n=48 total mice, n=24/group for preference and consumption. **p*<0.05 for correlation significance. All data are expressed as ±standard error mean.

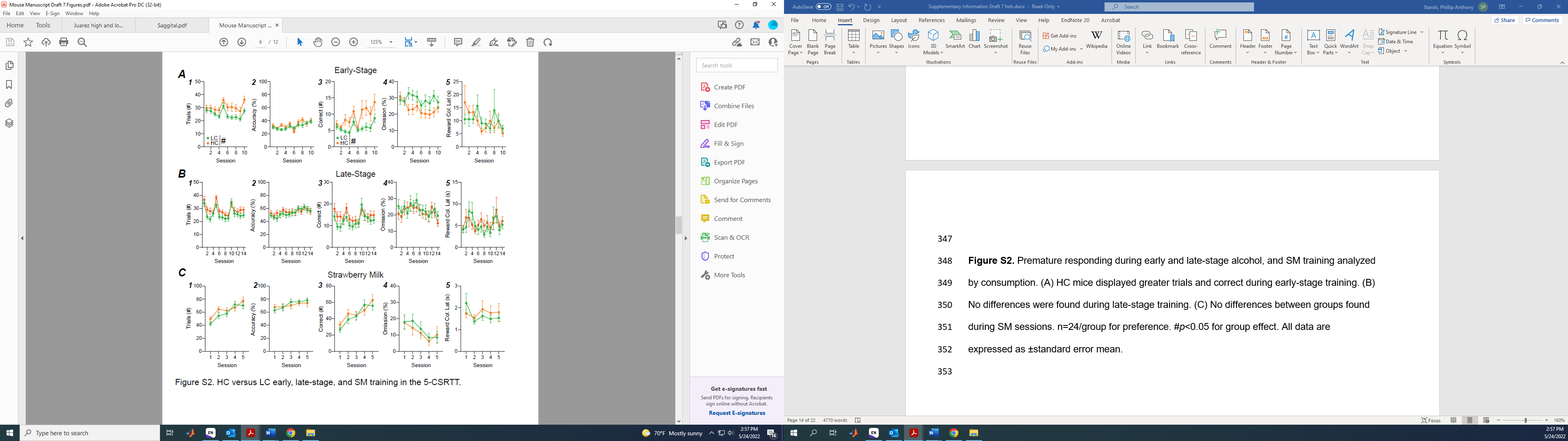

**Figure S2.** Premature responding during early and late-stage alcohol, and SM training analyzed by consumption. (A) HC mice displayed greater trials and correct during early-stage training. (B) No differences were found during late-stage training. (C) No differences between groups found during SM sessions. n=24/group for preference. #*p*<0.05 for group effect. All data are expressed as ±standard error mean.

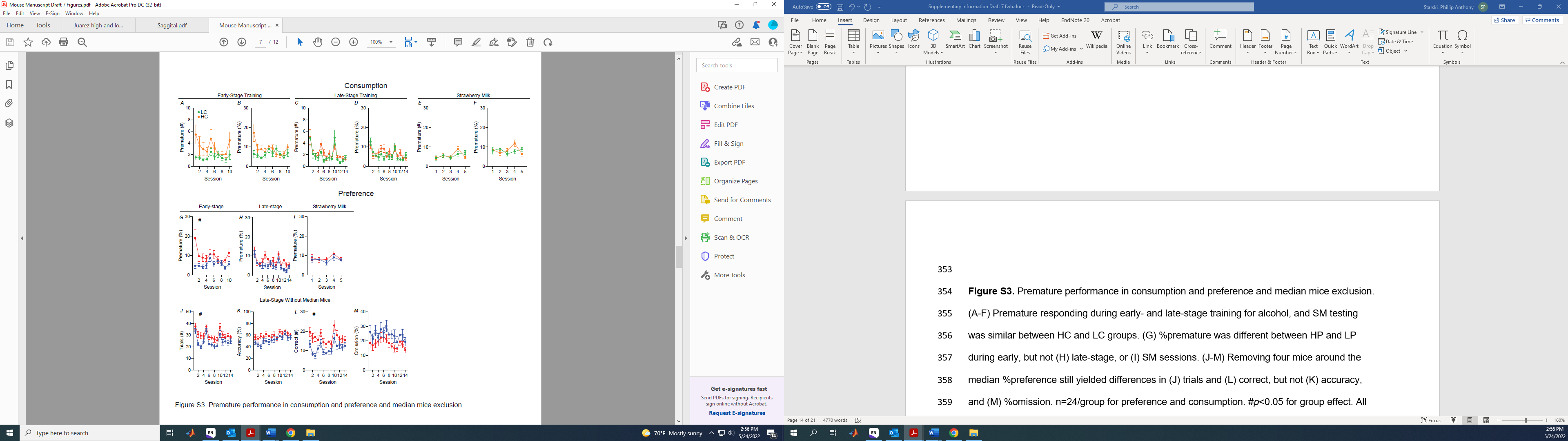

**Figure S3.** Premature performance in consumption and preference and median mice exclusion. (A-F) Premature responding during early- and late-stage training for alcohol, and SM testing was similar between HC and LC groups. (G) %premature was different between HP and LP during early, but not (H) late-stage, or (I) SM sessions. (J-M) Removing four mice around the median %preference still yielded differences in (J) trials and (L) correct, but not (K) accuracy, and (M) %omission. n=24/group for preference and consumption. #*p*<0.05 for group effect. All data are expressed as ±standard error mean.

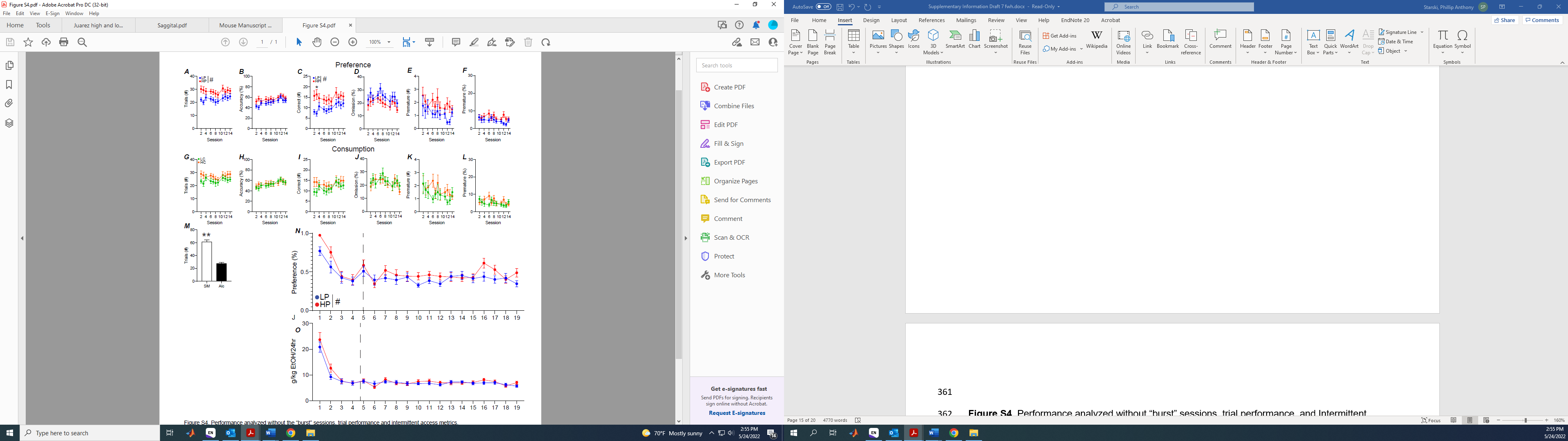

**Figure S4.** Performance analyzed without “burst” sessions, trial performance, and Intermittent Access metrics. Removal of the three “burst” sessions still showed differences in (A) trials and (C) correct during late-stage sessions, but not (B) accuracy, (D) %omission, (E) raw premature, and (F) %premature. (G-L) No differences were found by consumption in these measures. (M) Mice performed significantly more trials during SM sessions than alcohol. (N) %preference of HP and LP mice during IA2BC. (O) consumption of HP and LP mice during IA2BC. n=24/group for preference and consumption. #*p*<0.05 for group effect. **p*<0.05 for post-hoc significance. ***p*<0.01 for group effect. All data are expressed as ±standard error mean.

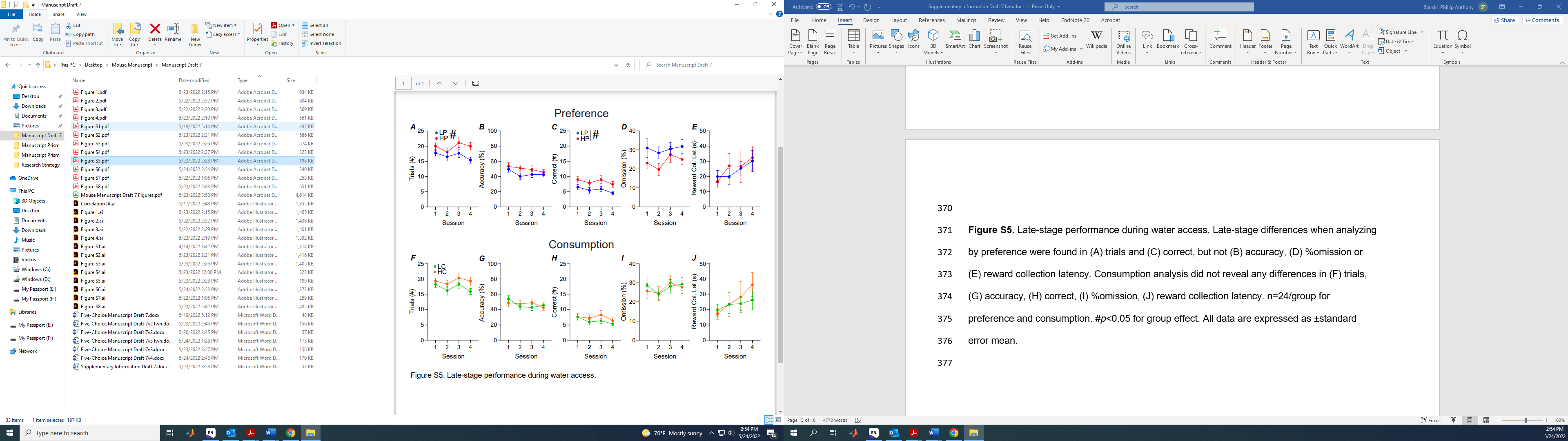

**Figure S5.** Late-stage performance during water access. Late-stage differences when analyzing by preference were found in (A) trials and (C) correct, but not (B) accuracy, (D) %omission or (E) reward collection latency. Consumption analysis did not reveal any differences in (F) trials, (G) accuracy, (H) correct, (I) %omission, (J) reward collection latency. n=24/group for preference and consumption. #*p*<0.05 for group effect. All data are expressed as ±standard error mean.

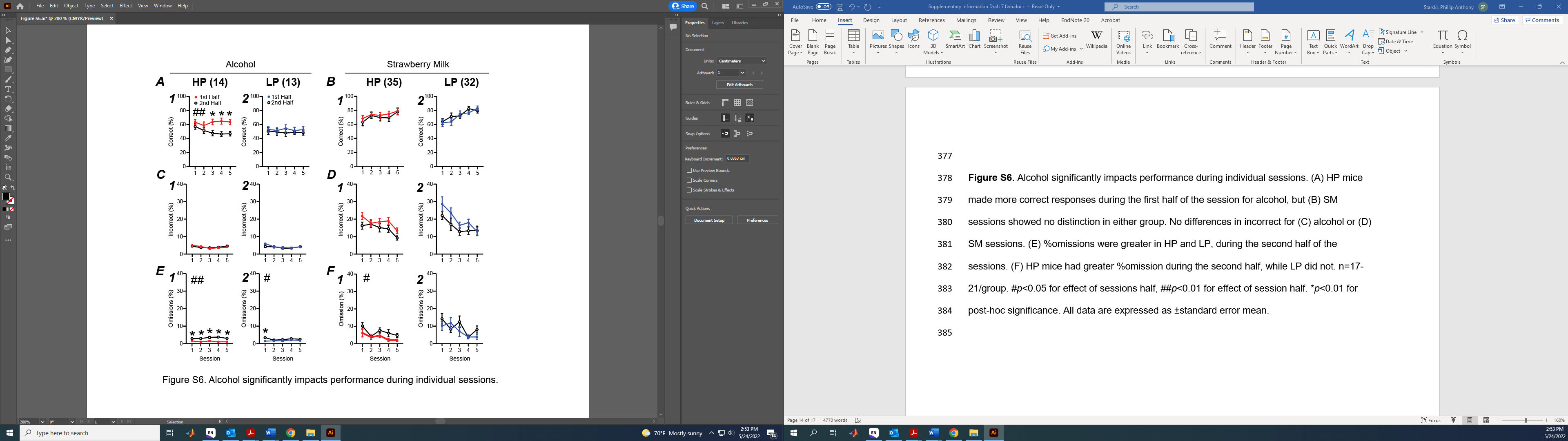

**Figure S6.** Alcohol significantly impacts performance during individual sessions. (A) HP mice made more correct responses during the first half of the session for alcohol, but (B) SM sessions showed no distinction in either group. No differences in incorrect for (C) alcohol or (D) SM sessions. (E) %omissions were greater in HP and LP, during the second half of the sessions. (F) HP mice had greater %omission during the second half, while LP did not. n=17-21/group. #*p*<0.05 for effect of sessions half, ##*p*<0.01 for effect of session half. **p*<0.01 for post-hoc significance. All data are expressed as ±standard error mean.

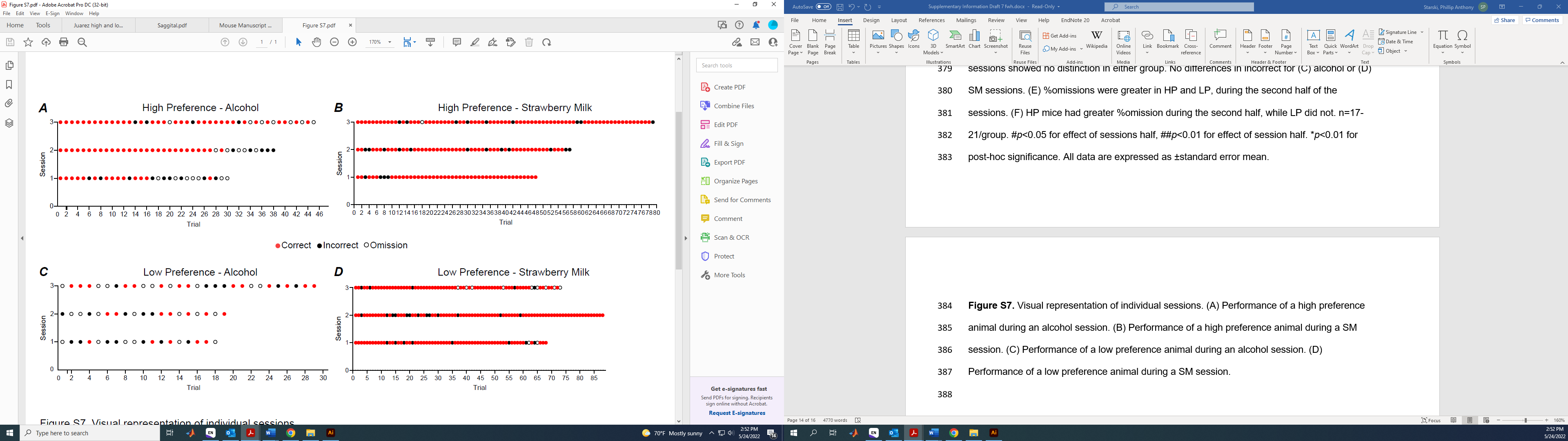
**Figure S7.** Visual representation of individual sessions. (A) Performance of a high preference animal during an alcohol session. (B) Performance of a high preference animal during a SM session. (C) Performance of a low preference animal during an alcohol session. (D) Performance of a low preference animal during a SM session.

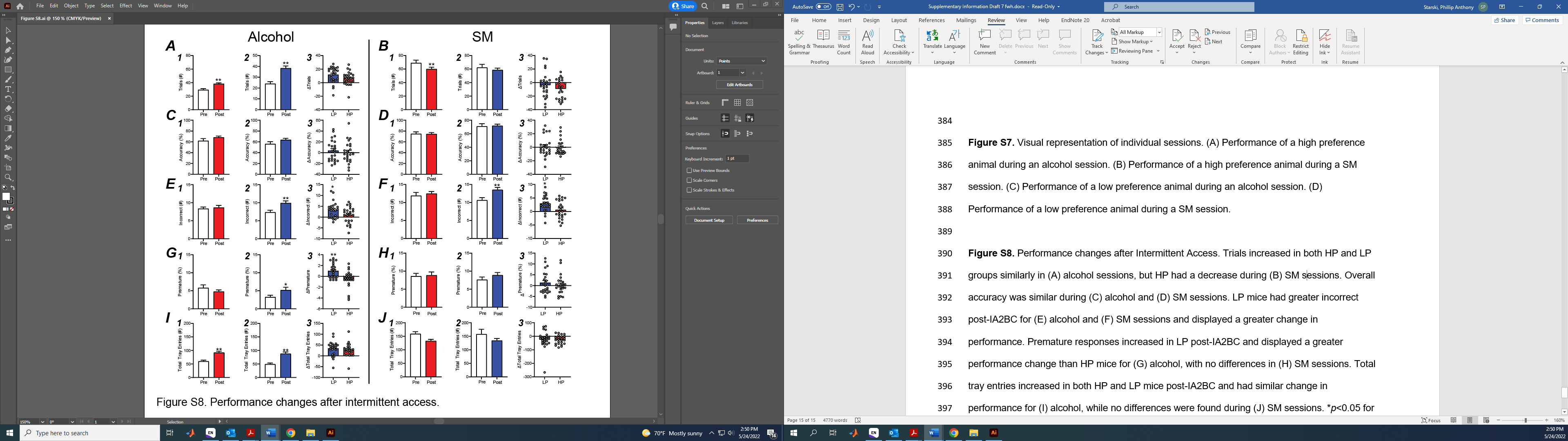

**Figure S8.** Performance changes after Intermittent Access. Trials increased in both HP and LP groups similarly in (A) alcohol sessions, but HP had a decrease during (B) SM sessions. Overall accuracy was similar during (C) alcohol and (D) SM sessions. LP mice had greater incorrect post-IA2BC for (E) alcohol and (F) SM sessions and displayed a greater change in performance. Premature responses increased in LP post-IA2BC and displayed a greater performance change than HP mice for (G) alcohol, with no differences in (H) SM sessions. Total tray entries increased in both HP and LP mice post-IA2BC and had similar change in performance for (I) alcohol, while no differences were found during (J) SM sessions. **p*<0.05 for group effect. ***p*<0.01 for group effect. All data are expressed as ±standard error mean.
